# Insulin resistance modifies longitudinal multi-omics responses to habitual diet

**DOI:** 10.64898/2026.02.17.706440

**Authors:** Heyjun Park, Xiaotao Shen, Dalia Perelman, Pranav Berry, Yingzhou Lu, Rachel Battersby, Sophia Miryam Schüssler-Fiorenza Rose, Alessandra Celli, Caroline Bejikian, Michael Snyder

**Affiliations:** Department of Genetics, Stanford University, Stanford, CA 94305; Department of International Health, Johns Hopkins Bloomberg School of Public Health, Baltimore, MD 21205; Singapore Phenome Center, Lee Kong Chian School of Medicine, Nanyang Technological University, Singapore, Singapore 639798; School of Chemistry, Chemical Engineering and Biotechnology, Nanyang Technological University, Singapore, Singapore 637371; Department of Medicine, Stanford University, Stanford, CA 94305; War Related Injury and Illness Study Center, Veteran Affairs Palo Alto Health Care System, Palo Alto, CA 94304

## Abstract

How habitual diet influences the gut microbiome and plasma metabolome across insulin resistance states remains unclear. We conducted year-long multi-omics profiling of 71 deeply phenotyped adults, integrating repeated assessments of diet, metabolome, gut microbiome, clinical laboratory measures, and inflammatory markers. Using gold-standard insulin suppression tests and machine learning-derived dietary patterns, we examined how dietary patterns relate to metabolic and microbial landscapes by insulin resistance status. Insulin-sensitive individuals exhibited stronger and more numerous diet-omics associations than insulin-resistant individuals, identifying metabolic flexibility as a central determinant of dietary responsiveness. Parabacteroides emerged as a candidate microbial mediator between refined carbohydrate-rich dietary patterns and host metabolic signatures. Integrated into a cardiovascular risk prediction model, diet, metabolites, microbial taxa, and immune markers each contributed to 10-year atherosclerotic cardiovascular disease risk. These findings show that inter-individual variation in cardiometabolic risk partly reflects differences in molecular responsiveness to habitual diet, informing precision nutrition and cardiovascular prevention.

## Introduction

Diet is a critical modifiable factor of metabolic health, acting not only through direct nutrient metabolism, but also through interactions with the gut microbiome and host physiological systems^1–3^. Diet shapes gut microbiome composition, leading to the production of bioactive metabolites, such as short-chain fatty acids (SCFA), bile acids, and branched-chain amino acids (BCAA), that regulate host metabolic pathways^4–7^. Multi-omics studies demonstrate that diet influences gut microbiome composition and host metabolic profiles, with these changes linked to metabolic health outcomes^8–13^.

In spite of the plethora of nutrition studies, the influence of habitual dietary patterns, as opposed to single nutrients or short-term diet, on microbiome composition and metabolomic profiles still remains poorly understood, especially in the context of metabolic disorders such as insulin resistance^14^. Insulin resistance is a major risk factor for cardiometabolic disorders, including type 2 diabetes, and has been associated with alterations in the gut microbial composition and systemic metabolism^14–16^. Much of the current existing research on diet-microbiome-metabolome interactions has been cross-sectional, limiting our understanding of how longitudinal dietary changes dynamically influence multi-omics profiles over time^3,9,10,17–19^. Nonetheless, long-term dietary habits appear to have a more profound effect on the gut microbial community^20^. Furthermore, it remains unclear whether insulin resistance alters the responsiveness of microbiome and metabolome to diet, highlighting the need for longitudinal analyses to better understand these interactions and ultimately inform targeted dietary interventions tailored to an individual’s metabolic capabilities.

Recent advances in multi-omics profiling and machine learning have provided new opportunities to investigate the complex relationships between metabolism, the microbiome and health^9,10,18,19,21^. Beyond assessing nutrients, holistic dietary patterns, including those identified through data-driven approaches, offer new insights into habitual diet and its metabolic consequences^13,22–24^. However, the extent to which dietary patterns shape metabolome-gut microbiome relationships, and the degree to which microbial taxa mediate diet-metabolome associations through specific biological mechanisms are not yet fully established. Additionally, integrating longitudinal dietary changes with multi-omics profiling allows for a deeper understanding of the temporal complexity of diet-ome interactions. In addition, few studies have explored how insulin resistance status modifies these dynamics. Therefore, research is needed to incorporate both dietary complexity and individual metabolic variability.

To address these gaps, we leveraged a longitudinal multi-omics study in a well-characterized cohort of generally healthy adults with repeated habitual dietary assessments and high-resolution profiling of the metabolome, gut microbiome, and clinical laboratory markers. To overcome cross-sectional limitations and capture temporal dynamics, we employed longitudinal sampling at multiple time points. To move beyond single nutrient analyses, we applied machine learning to derive holistic dietary patterns and systematically examined how these patterns shape molecular and microbial landscapes. To test whether insulin resistance modified diet-microbiome-metabolome associations, we stratified analyses by insulin resistance status and applied mediation analyses to identify microbial taxa mediating these relationships. As a translational application of these insights, we integrated multiple data types (diet, omics, clinical labs, inflammatory markers, and demographics) to predict 10-year atherosclerotic cardiovascular disease (ASCVD) risk and assess the relative importance of each data modality to disease prediction. Our findings reveal the dynamic nature of diet-ome interactions and demonstrate how habitual diet influences metabolic health across insulin resistance states. This work informs precision nutrition strategies that account for individual metabolic status for disease prevention.

## Results

### Study Design and Cohort Characteristics

We have previously established a cohort of >100 adults at risk for type 2 diabetes (T2D) and longitudinally followed them between 2013-2017^25–28^. In this cohort, we collected blood and stool samples every 3 months, and the participants completed a detailed dietary assessment. Through longitudinal multi-omics profiling, we deeply phenotyped the participants at each visit. Metabolites, cytokines, and clinical laboratory markers were measured from plasma samples, and gut microbes were analyzed from stool samples. Specifically, for the metabolite detection, liquid chromatography with tandem mass spectrometry (LC-MS/MS) was employed for untargeted metabolomics analysis, and for gut microbe detection, 16S ribosomal RNA gene sequencing was employed for microbiome analysis (Methods)^25–27^. The study was approved by the Stanford University Institutional Review Board (IRB #23602), and all study participants consented.

An overview of the study design is presented in **Figure 1A**. For this study, we analyzed data from 71 participants (mean age 53.0+7.6 years; 56% female; BMI 28.6+3.8; **Figure 1B**) who provided a food diary for at least one day during their quarterly visits. These participants were followed across four time points over an average period of 335 days. The average time between each of the two data collection points was approximately three months. This ensured identifying meaningful longitudinal dietary trends. During each visit, a one-day food diary was provided by participants, capturing a day representative of their typical intake during the previous three months.

**Figure 1.**
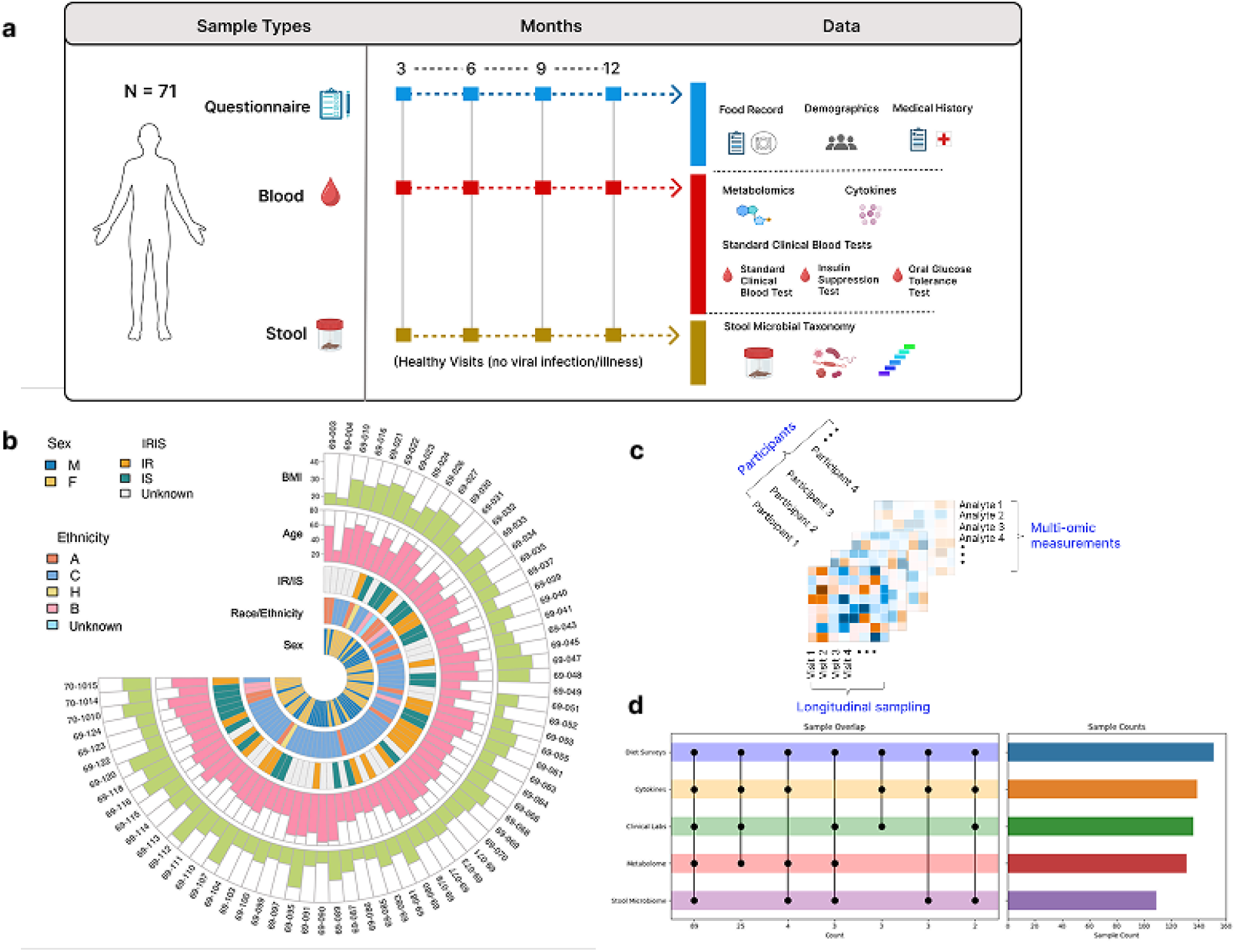
Study design, participant characteristics, and multi-omics data structure. **a**, Study design. Participants completed quarterly visits over one year that included food records, demographic and medical history questionnaires, clinical laboratory tests, and multi-omics profiling. At baseline, participants underwent standardized metabolic tests, including an insulin suppression test (IST) and an oral glucose tolerance test (OGTT). IST determined participants’ insulin resistant status. **b,** Baseline characteristics. Circular plot depicting each participant’s sex, insulin resistant or insulin sensitive status, race/ethnicity, age, and body mass index. Inner layers denote categorical variables, and outer layers display continuous measures. **c,** Longitudinal data collection. Diagram showing repeated multi-omics measurements collected from each participant across multiple study visits. d, Data availability and overlap across data types. Sample counts for each data type (diet surveys, cytokines, clinical laboratory assays, plasma metabolomics, and stool microbiome) and the extent of overlap across these data layers. Illustration in a created with BioRender.com.

Dietary data were matched with corresponding omics data collection dates to investigate potential relationships between changes in diet and omics profiles (**Figure 1C**, **Methods**). From quarterly visits, we obtained 151 food diaries, 139 plasma samples, and 109 stool samples. These were used to generate the comprehensive dataset for each participant that included annotated metabolites (n = 724), cytokines, chemokines, and growth factors (n = 62), clinical labs (n = 43) and gut microbes (multiple taxonomic levels; n = 109). The dietary data included nutrient (n = 62) and food group variables (n = 20) (**Figure 1D, Methods**).

In addition to multi-omics profiling, this cohort underwent a thorough evaluation of glucose control capacity, utilizing standard metrics such as fasting plasma glucose (FPG) and Hemoglobin A1c (HbA1c). 45 participants underwent more in-depth assessments such as an annual oral glucose tolerance test (OGTT) and an insulin suppression test (IST) (**Figure 1A**). Based on the IST results, which measure steady-state plasma glucose levels (SSPG), participants were categorized as either insulin-sensitive (IS; SSPG<150 mg/dL; n = 18), or insulin-resistant (IR; SSPG≥150 mg/dL; n = 27). The baseline characteristics related to glucose metabolism among IS and IR are also provided in **Table 1**. By leveraging this extensive and rich longitudinal profiling dataset of the cohort, this study examined the relationships between longitudinal diet and omics signatures in individuals at risk for T2D. Additionally, this study also explored the influence of insulin resistance on these relationships.

**Table 1.**
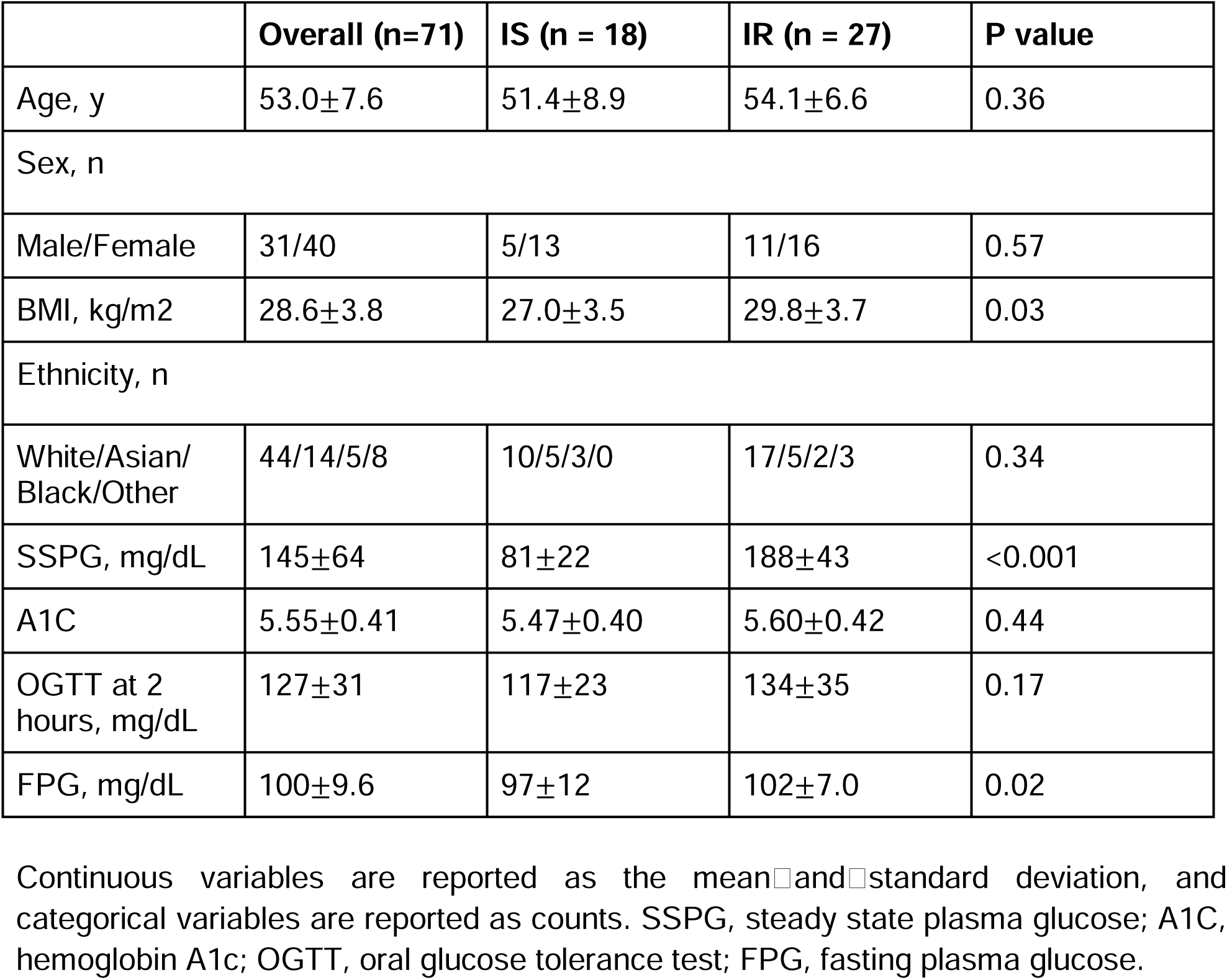
Baseline Characteristics.

### Longitudinal dietary changes in the study participants

We identified significant longitudinal changes in habitual diet among participants (n = 71) from baseline (T1) to endline (T4). Using linear mixed effect models with baseline as the reference, 13 nutrients showed significant changes over time in the overall cohort based on *P_adj_* < 0.2 (**Figure 2A**). For example, total fiber, choline, and biotin intakes increased over the course of the study. Vitamin B3 and fat-soluble vitamins (alpha-tocopherol, and vitamin K), and potassium also increased from T1 to T4. Changes were also observed in 6 food categories in the overall cohort, such that intakes of non-starchy vegetables, eggs, and meats and poultry increased over the course of the study. Consumption of rice, pasta, and noodles increased from the beginning to T2, and tea intake also increased from T1 to T3. In contrast, intake of mixed meal items such as pizza, sandwiches, and wraps declined through the study.

**Figure 2.**
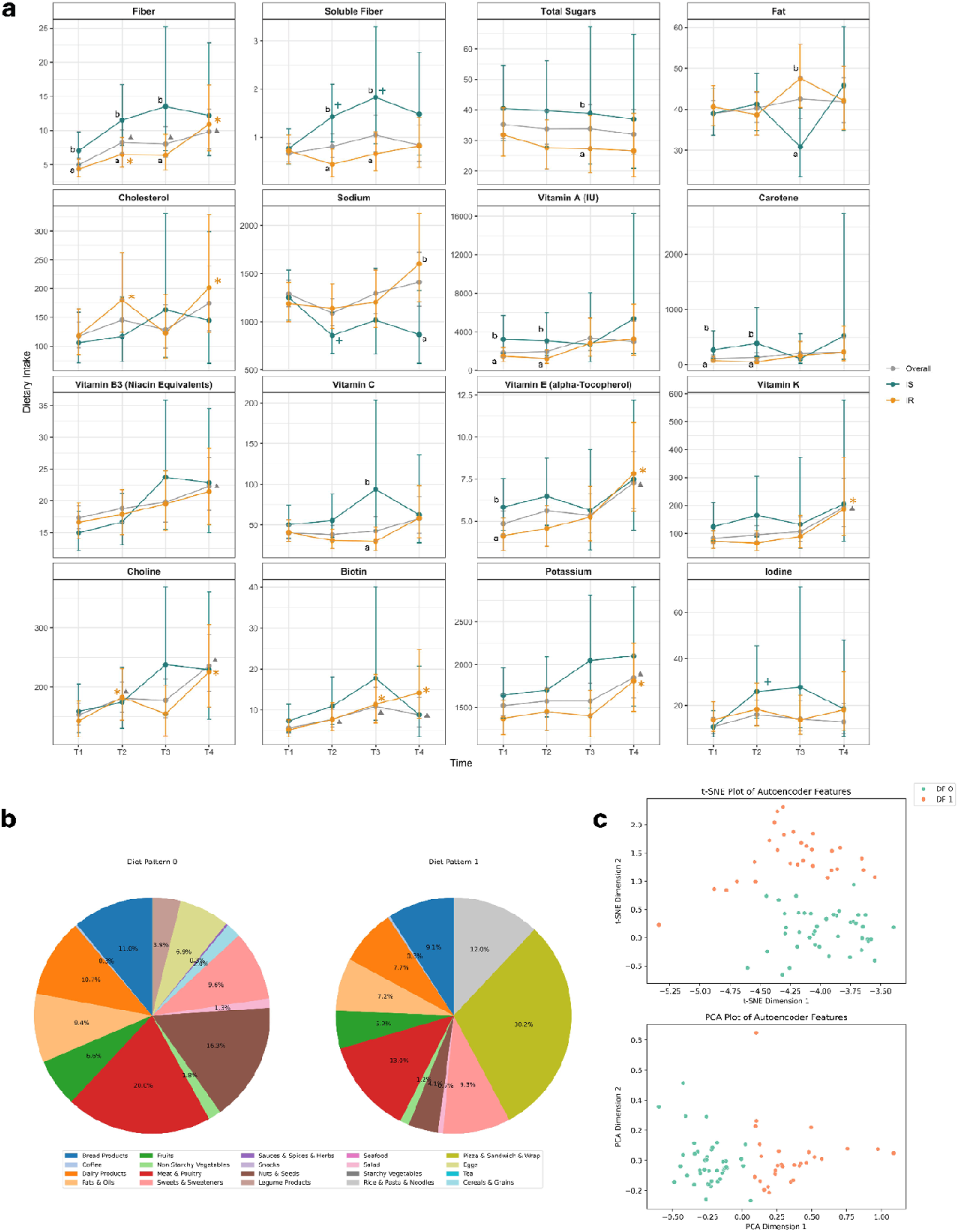
a, Longitudinal changes in nutrient intake over one year in the overall cohort and by insulin resistance status. Predicted mean intakes and 95% confidence intervals are shown for the overall cohort (n=71;grey), insulin-sensitive (IS; n=18; green) and insulin-resistant (IR; n=27; orange) participants. Estimates were obtained from linear mixed-effects models with baseline (T1) as the reference. Significant changes from baseline are indicated by symbols (triangle: overall cohort; cross: IS; asterisk: IR; *P_adj_* < 0.20). Differences between IS and IR at each time point are denoted by distinct letters (*P_adj_* < 0.20) based on Mann-Whitney tests. Among the 25 nutrients that showed at least one significant within-group change or IS-IR difference, 16 representative nutrients are displayed. **b,** Energy distribution across food groups for each dietary pattern cluster (DP0 and DP1). Each slice represents the energy contribution (%) of a given food group, with colors indicating distinct food categories. DP1 shows higher intakes of refined carbohydrate sources (pizza, sandwiches, tacos, wraps, rice, pasta, and noodles) than DP0, therefore, represents a pattern that is richer in refined carbohydrate. **c,** Visualization of latent dietary features derived from the autoencoder. Top: t-SNE projection of the latent feature space. Bottom: PCA projection of the same features. Each point represents an individual participant, colored by assigned dietary pattern cluster (DP0 or DP1). The clear separation between clusters in both embeddings demonstrates that the latent features learned by the autoencoder represent distinct and coherent dietary patterns.

### Differences in dietary intakes between individuals with IR and IS

Given the well-characterized insulin resistance phenotypes in our cohort (n=45), we assessed dietary intakes separately within insulin-sensitive (IS) and insulin-resistant (IR) participants using linear mixed-effect models with baseline (T1) as the reference, and compared dietary differences at each time point using Mann-Whitney tests (*P_adj_* < 0.2; **Figure 2A**). Several nutrients differed between IS and IR participants across the study period. Both groups showed increases in total fiber and soluble fiber, although IS participants maintained consistently higher levels through the study. Total sugar and vitamin C intakes were also generally higher in IS group, with significant differences at T3. Vitamin A, carotene, and vitamin E were significantly higher in IS individuals at the beginning of the study (T1 and T2). In contrast, fat intake decreased in IS group and reached significantly lower levels than IR at T3, and sodium showed a transient reduction in IS at T2 and remained significantly lower than IR at T4. Within the IR group, dietary cholesterol increased at T2 and T4 relative to baseline, and fat-soluble vitamins (vitamin E and vitamin K), biotin and choline showed significant increases over the study period. Iodine intake increased in IS group, whereas potassium increased in the IR. Together, these results highlight distinct dietary trajectories between IS and IR participants over the one-year follow-up.

### Machine learning identified two distinct habitual dietary patterns in the cohort

Dietary intake patterns in the cohort were highly complex, influenced by numerous dietary components that varied across individuals over time. To systematically identify these patterns, we applied an autoencoder analysis, a neural network for nonlinear dimensionality reduction. This method captures latent dietary structures with minimal reconstruction error. Autoencoder modeling followed by k-means clustering was performed using intakes of energy, carbohydrates, and food weight data across food groups over the study period. After extensive validation, including the evaluation of 1,140 autoencoder models, two clusters of individuals, Dietary Pattern 0 (DP0; n=42 participants) and Dietary Pattern 1 (DP1; n=29 participants), were identified with high clustering stability and strong reproducibility (**Figures 2B**, and **Methods**). These patterns were further confirmed using principal component analysis (PCA) and t-distributed stochastic neighbor embedding (t-SNE), both of which clearly separated participants by these dietary classifications (**Figure 2C**).

At the participant level, DP0 was characterized by significantly higher median energy proportions from meat and poultry (12%), nuts and seeds (11%), and bread products (8%) compared with DP1 (Mann Whitney tests with BH correction; *Padj* < 0.20). In contrast, DP1 showed significantly higher median energy proportions from sandwiches, wraps, tacos, and pizza (16%), as well as rice, pasta, and noodles (7%). When summarized at the pattern level, these distinct food groups comprised over 45% of total caloric intake in DP0 and over 40% in DP1, respectively (**Figure 2B**). Subsequent nutrient comparisons revealed that fiber (both total and soluble), vitamin K, copper, magnesium, and potassium were significantly lower in DP1, while sodium levels were higher in DP1 than DP0 (*P*_adj_ < 0.2). Taken together, these characteristics indicate that, compared to DP0, DP1 reflects a dietary pattern that is lower in fiber and richer in refined carbohydrates.

### Cross-sectional correlations among habitual diet, plasma metabolome, and gut microbiome in the overall cohort

We assessed cross-sectional associations between 62 nutrients and 724 plasma metabolites (**Methods**). Many energy-adjusted nutrients and food groups showed correlations with metabolites (*P_adj_* < 0.20, all correlations with an absolute value greater than 0.4; n=81 correlations), even after adjusting for age and sex (**Figure 3A**). Pathway enrichment analyses using KEGG databases (**Methods**) identified the top 10 metabolic pathways, including glutathione metabolism, histidine metabolism, and monoacylglycerol, significantly related to 7 nutrient groups (i.e., carbohydrates, protein, fat, water-soluble vitamins, fat-soluble vitamins, minerals, caffeine; P < 0.01; **Figure 3B**). When the pathways were grouped by broader chemical classifications, over 45% were involved in amino acid metabolism, followed by lipid metabolism (21%) and xenobiotic metabolism (12%; **Figure 3C**). Interestingly, about 25% of the identified metabolites were phospholipids (i.e., phosphatidylethanolamine (PE) and phosphatidylcholine (PC)), which were significantly associated with various nutrients such as carbohydrates, protein, cholesterol, choline, and selenium. However, these phospholipids did not have specific annotations for distinct metabolic pathways.

**Figure 3.**
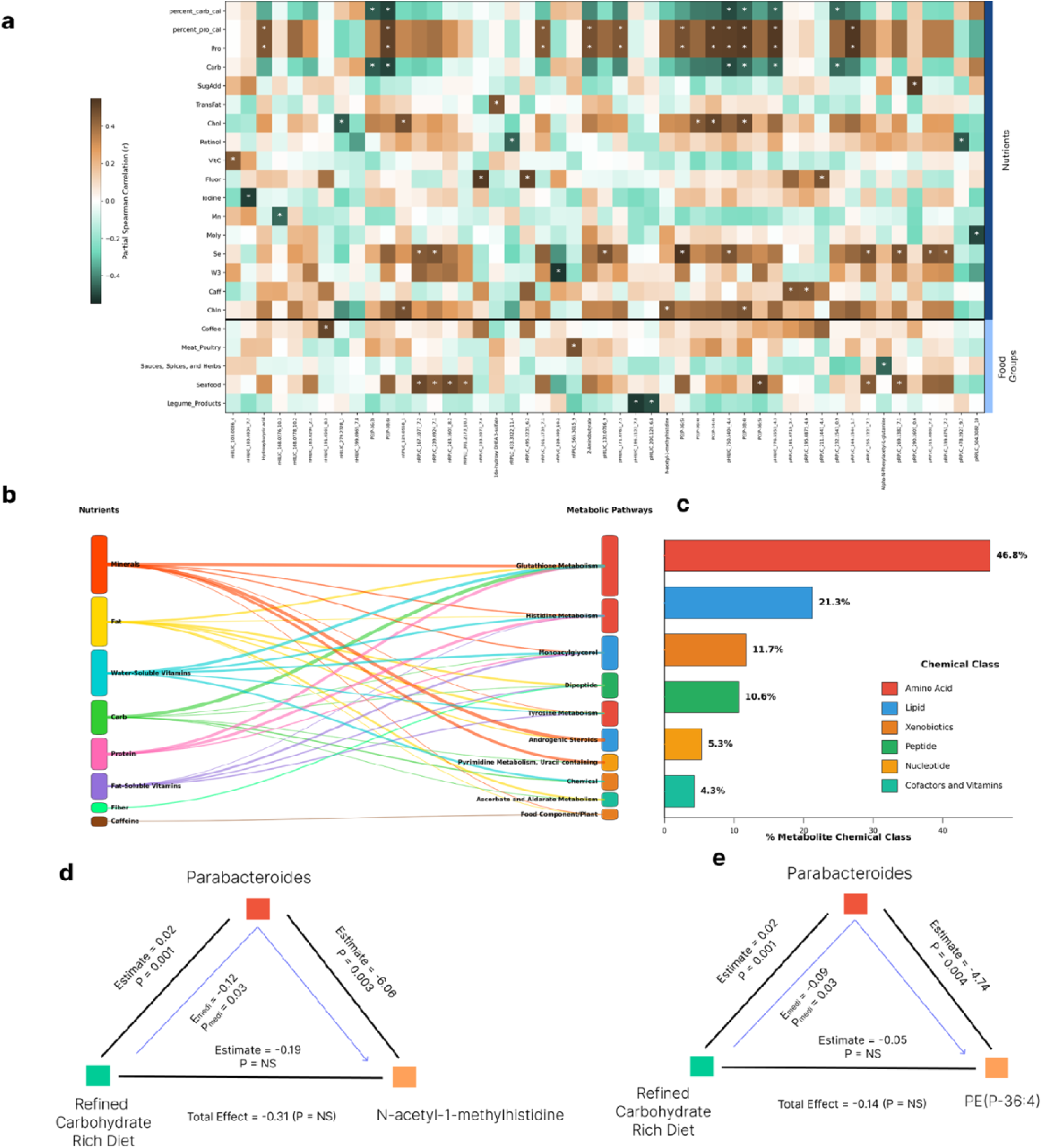
Cross-sectional correlations and mediation analyses linking habitual diet, plasma metabolome, and the gut microbiome in the study cohort. **a**, Partial Spearman correlations between habitual dietary components (nutrients and food groups) and plasma metabolites, adjusted for age and sex. Asterisks denote significant associations (*P_adj_* < 0.20). **b,** Sankey diagram connecting nutrient groups to the top enriched metabolic pathways. Pathway enrichment was conducted using KEGG annotations and highlights pathways significantly associated with seven nutrient groups. **c,** Distribution of enriched pathways across broader metabolite chemical classes. **d,** Mediation of the association between refined carbohydrate diet and N-acetyl-1-methylhistidine via Parabacteroides. The blue arrow shows the significant indirect (mediated) effect (DP1→Parabacteroides→ N-acetyl-1-methylhistidine). The direct effect of diet on N-acetyl-1-methylhistidine, not passing through Parabacteroides, was not significant, indicating full mediation. **e,** Mediation of the association between refined carbohydrate diet and PE(P-36:4) via Parabacteroides. The blue arrow shows the significant indirect (mediated) effect (DP1→ Parabacteroides→PE(P-36:4)). The direct effect of diet on PE(P-36:4) was not significant, indicating full mediation.

Energy-adjusted dietary components also showed cross-sectional associations with gut microbiota after adjustment for age and sex (*P* < 0.05). Nutrients most strongly correlated with microbial taxa included fiber, vitamin E, pantothenic acid, monounsaturated fat, omega-6 fatty acids, and monosaccharides. Eight genera (*Anaerotruncus, Parabacteroides, Bilophila, Butyricimonas, Clostridium sensu stricto, Clostridium XlVb, Lachnospiracea incertae sedis, and Roseburia*) were associated with at least seven dietary components. *Butyricimonas, Clostridium XIVb*, and *Roseburia*, have previously been linked to high fiber diets^29–34^, and are known to produce short-chain fatty acid (SCFA)^35–39^. *Erysipelotrichaceae incertae sedis*, previously related to dietary fat and host lipid metabolism^40,41^, was associated with fat-soluble vitamin A in our cohort. Coffee intake was positively correlated with *Butyricimonas*, though the biological mechanism underlying this relationship remains unclear. In addition, machine learning (ML)-derived longitudinal dietary patterns were significantly associated with 4 microbial taxa (*Parabacteroides, Anaerotruncus, and Flavonifractor* at genus levels and unclassified *Porphyromonadaceae*). However, these associations did not remain statistically significant after further multiple testing adjustment.

### Microbiome-mediated effects in the connections from dietary patterns to molecular profiles in the overall cohort

We next explored potential causal connections between dietary components, gut microbial taxa, and host metabolites through mediation analysis. This approach assesses whether the significant microbial taxa identified earlier mediated any of the associations between the dietary components and plasma metabolites (**Figure 3A**). Our analysis revealed 6 significant directional mediation effects (*P_medi_* < 0.05) involving the 2 microbial taxa from diet to the 6 metabolites in 3 known metabolic pathways (histidine metabolism, plasmalogen, and acetylated peptides metabolism). Notably, the genus *Parabacteroides* significantly mediates the relationship between the refined carbohydrate rich pattern (DP1) and the metabolites N-acetyl-1-methylhistidine and PE(P-36:4) (indirect effect estimates -0.12 and -0.09, respectively; **Figure 3D, 3E)**. These findings suggest that refined carb-richer dietary pattern significantly modifies certain microbial taxa, and these microbial changes in turn mediate shifts in host molecular profiles, potentially leading to significant physiological outcomes.

### Differential relationships between longitudinal changes in nutrient intakes and plasma metabolome by insulin-resistant status

After identifying cross-sectional associations between average dietary intakes and omics profiles, we next shifted our focus to the longitudinal associations using a multidimensional feature-based approach that captures the dynamic nature of nutrient intakes and corresponding molecular changes over time. Specifically, we examined whether insulin-sensitive (IS) and insulin-resistant (IR) individuals demonstrate distinct metabolic responses to dietary changes.

We represented the longitudinal changes for each nutrient using 11 features that summarized temporal intake dynamics across 4 time points (early and late slopes, cumulative exposure (AUC), and peak changes; **Figure 4A; Methods)**. We then correlated them with corresponding features extracted from the metabolome data. While both IS and IR groups had the same number of metabolites correlated to dietary changes (n = 169), the IS group exhibited more total significant correlations compared to the IR group (1,777 correlations vs. 1,571; both q < 0.05) and stronger correlations (mean |r| = 0.67 vs. 0.41; *P* < 2.2e-16; Wilcoxon test; **Figure 4B-4D**). Pathway enrichment analyses using KEGG databases on the metabolites showed that IS individuals had 6 enriched pathways, such as primary bile acid biosynthesis, caffeine metabolism, phenylalanine metabolism, steroid hormone biosynthesis, sphingolipid metabolism, and pyrimidine metabolism (*P* < 0.05). In contrast, IR individuals showed enrichment in only 2 metabolic pathways such as bile acid biosynthesis and caffeine metabolism (*P* < 0.0005).

**Figure 4.**
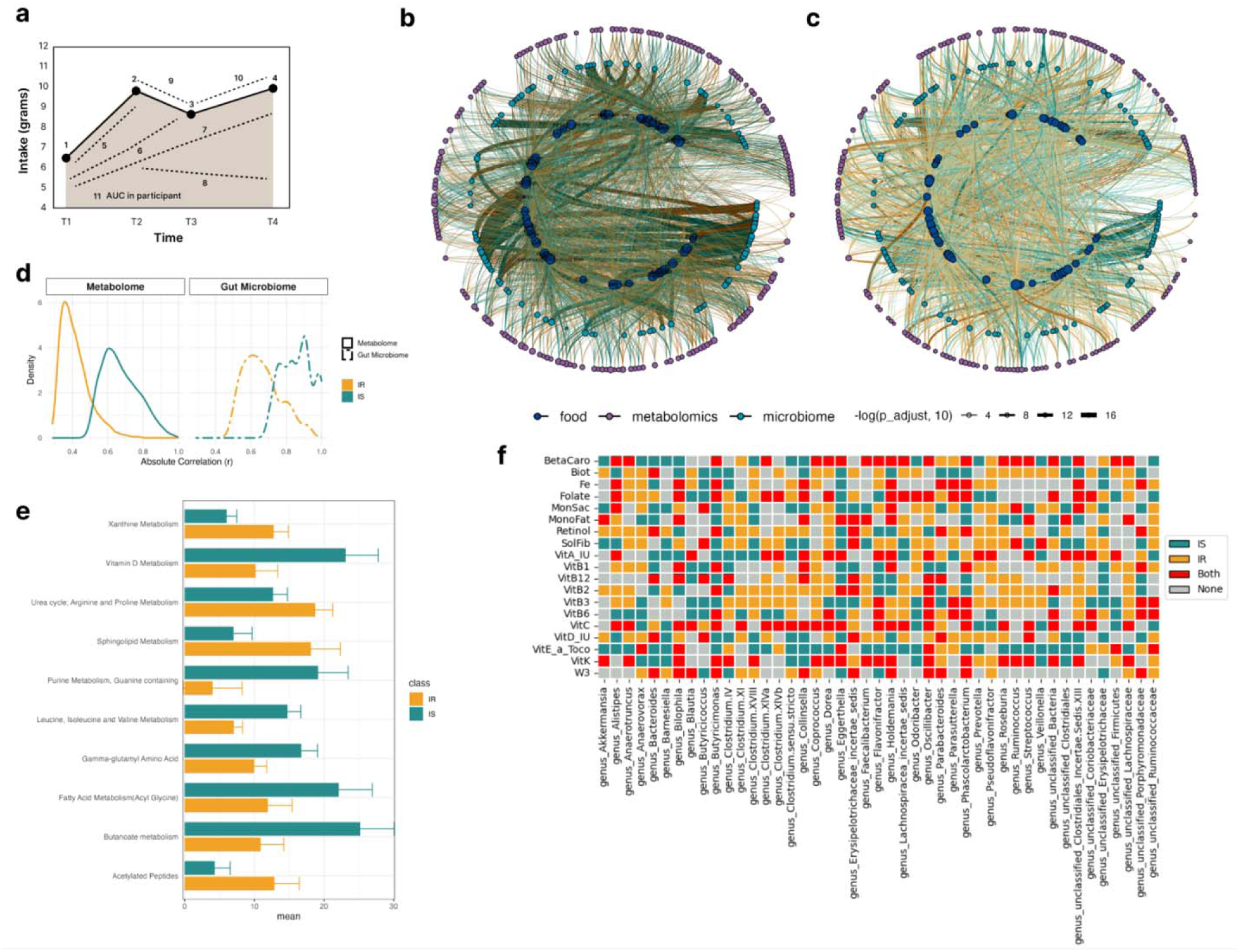
Longitudinal nutrient dynamics and their multi-omics associations differ by insulin-resistance status. **a**, Illustration of the longitudinal change metrics to summarize diet and omics trajectories across study time points. 11 temporal features including early and late slopes, cumulative exposure (area under the curve), and peak change, were extracted for each analyte in each participant. **b,** Significant longitudinal associations between change features of nutrient and ones of multi-omics in insulin-sensitive (IS) individuals (q<0.05). Nodes are arranged in three concentric layers: nutrients (inner layer), gut microbial taxa (middle layer), and plasma metabolites (outer layer). Edges represent significant correlations between temporal change features across omics layers. **c,** Significant longitudinal associations between change features of nutrient and ones of multi-omics in insulin-resistant (IR) individuals (q<0.05). As in panel b, the inner layer indicates nutrients, the middle layer represents gut microbial taxa, and the outer layer represents plasma metabolites. **d,** Distribution of absolute correlation strengths between change features of nutrient and omics in IS (green) and IR (orange) individuals. Kernel density estimates are shown separately for metabolomic associations (left) and gut microbiome associations (right). Only correlations with adjusted p values < 0.05 are shown. **e,** Pathway-level comparison of the nutrient-metabolite association density between IS and IR individuals. For each enriched pathway (q<0.05), density was calculated by normalizing the total number of significant nutrient-metabolite correlations by the number of distinct metabolites mapped to that pathway. The top ten pathways by density are shown. **f,** Nutrients driving longitudinal nutrient-microbiome associations in IS and IR individuals. The heatmap summarizes significant longitudinal associations between change features of nutrient and genus-level gut microbial taxa (q<0.05) in IS and IR individuals. Each cell indicates whether a given nutrient-genus association was significant in IS only (green), IR only (orange), both groups (red), or neither group (gray). Rows indicate individual nutrient intakes, and columns represent gut microbial genera.

To further compare the pathway-level metabolic responses between IS and IR individuals, we quantified the density of nutrient-metabolite associations within each sub-pathway. For each group, the density was calculated as the total number of significant associations (q < 0.05) divided by the number of distinct metabolites mapped to that sub-pathway (**Methods**). Then, we applied a bootstrapped permutation to repeatedly resample the data and test for differences in pathway density between the groups using Wilcoxon rank-sum tests. This analysis showed clear pathway-level differences in the metabolic response to dietary change. IS individuals showed significantly greater density of associations in vitamin D metabolism, purine metabolism, branched-chain amino acid metabolism and gamma-glutamyl amino acid pathways (*P* < 0.05; **Figure 4E**). In contrast, IR individuals exhibited greater density in acetylated peptides, sphingolipid metabolism, and xanthine metabolism. Together, these findings demonstrate that IS and IR individuals differ not only in the overall strength of nutrient-metabolite associations but also in which metabolic pathways show the highest per-metabolite responsiveness to dietary changes.

### Varied relationships between longitudinal changes in nutrient intakes and microbiome by insulin-resistant status

Using the same feature-based approach applied to nutrient dynamics, we explored associations between longitudinal changes in nutrient intakes and gut microbial taxa by IS/IR status. The number of significantly correlated gut microbial taxa was similar between groups (94 taxa in IS and 96 in IR). However, similar to our findings in metabolome, both the magnitude and frequency of these correlations differed. IS individuals showed a greater total number of significant correlations than the IR (1,164 vs. 1,100; q-val<0.05 for both; **Figure 4B, 4C)** and stronger associations (mean |r| = 0.86 vs. 0.67; p value < 2.2e-16, Wilcoxon rank sum test; **Figure 4D**).

Next, we identified the specific gut microbial genera that most accounted for these associations. In individuals with IS, the genus *Coprococcus* significantly correlated with changes in 22 nutrients (q<0.05; **Figure 4B**). Among IR individuals, the genera *Ruminococcus* and *Butyricicoccus* each significantly correlated with changes in 20 nutrients (q < 0.05; **Figure 4C**). To complement these findings and identify the dietary components driving these associations, we subsequently examined the specific nutrients that predominantly contributed to these associations in each IS/IR group (**Figure 4F**). In IS participants, intake changes in monosaccharides, vitamin D, iron, biotin, manganese, and soluble fiber were significantly correlated with 31 microbial genera out of total 43 genera (q < 0.05), accounting for 19% of all significant genus level correlations (n = 109). In IR individuals, changes in vitamin C, folate, phosphorus, omega-3 fatty acid, iron, and added sugar were significantly correlated with 37 microbial genera out of total 45 genera (q < 0.05), accounting for 22% of all significant genus level correlations (n = 114).

Our feature-based approach also detected dynamic, temporal relationships between nutrients and gut microbes **(Methods)**. We observed both lagged and synchronized associations. Specifically, early increases in manganese intake were followed by later rises in the abundances of *Clostridium sensu stricto* and *Flavonifractor*, whereas those same initial manganese increases preceded decreases in *Parasutterella* and *Veillonella*. Similarly, higher alcohol intake at earlier time points was associated with a subsequent increase in *Eggerthella*. In addition, we found synchronous relationships for several micronutrients. Vitamin K intake rose and fell in parallel with *Parasutterella* abundance, and changes in Vitamin A and carotene did with *Holdemania*. Manganese also fluctuated together with *Clostridium sensu stricto*. These findings demonstrate how our longitudinal, shape-feature approach captures both immediate (synchronized) and lagged diet-microbiome interactions.

### Dietary patterns and clinical and inflammatory response markers in insulin resistance

We examined how insulin resistance (IR) status is associated with clinical laboratory markers in the context of dietary patterns (i.e., IS-DP0 (n = 13), IS-DP1 (n = 5), IR-DP0 (n = 16), and IR-DP1 (n = 11)). At each visit, 43 clinical laboratory tests and 62 cytokine analyses were performed (**Methods**), revealing diet-dependent associations between insulin resistance and circulating cytokines, as evidenced by significant interactions between SSPG and dietary pattern for several cytokines including leptin and VEGFD (*P_adj_* < 0.02; **Figure 5A**). For example, at higher SSPG levels (indicating greater IR), participants showing DP1 (the refined carbohydrate rich dietary pattern) showed lower leptin concentrations, whereas those on DP0 showed relatively higher leptin levels across the SSPG range. These results were observed after adjustment for age, sex, race/ethnicity, and BMI, suggesting that a dietary pattern, specifically a refined carbohydrate rich diet, may influence leptin regulation among IR individuals independent of one’s overall adiposity. In contrast, at higher SSPG levels, participants showing DP1 exhibited lower concentrations of VEGFD, while this pattern was attenuated among those on DP0. This finding is consistent with diet-related differences in angiogenic or vascular remodeling in the IR subgroup (**Figure 5A**).

**Figure 5.**
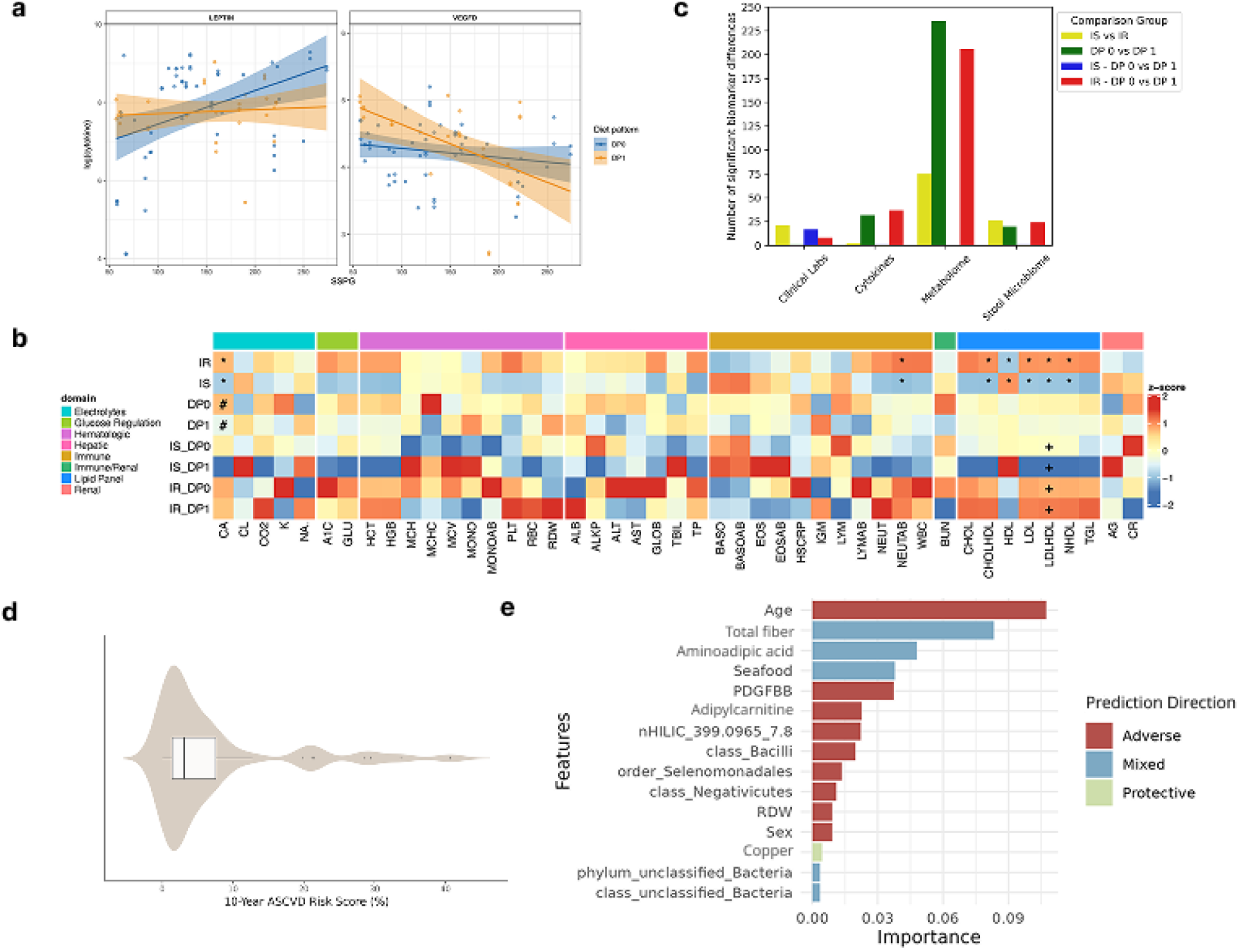
Biomarker heterogeneity associated with insulin resistance and dietary patterns across multi-omics layers and cardiovascular risk. Dietary pattern-dependent heterogeneity of insulin-resistance-associated biomarker profiles across multi-omics layers and cardiovascular risk **a,** Relationships between steady-state plasma glucose (SSPG) and log-transformed circulating leptin and VEGFD, stratified by dietary pattern (DP0, blue; DP1, orange). Lines indicate fitted values from linear mixed-effects models with 95% confidence intervals, adjustment for age, sex, race/ethnicity, and baseline BMI. Interaction effects between SSPG and dietary pattern were identified (*P_adj_* < 0.2) **b,** Heatmap of differences in clinical laboratory markers across insulin resistance status (IS, IR) and diet pattern groups (DP0, DP1). Rows also represent the four subgroup combinations (IS-DP0, IS-DP1, IR-DP0, IR-DP1). Values are standardized estimated marginal means (z-scores) from linear mixed-effects models. Blue indicates lower concentrations and red indicates higher values for each biomarker. Symbols denote BH-adjusted significance of model terms: * indicates a main effect of insulin resistance, # indicates a main effect of diet pattern, and + indicates a significant interaction between insulin resistance and diet pattern (*P_adj_* < 0.20). For biomarkers with significant interactions, Tukey post-hoc tests were performed among the four subgroups. Biomarkers are grouped by physiological domain. **c,** Differential biomarker associations with insulin resistance and dietary pattern across multi-omics layers. Bar plots summarize the number of biomarkers with significant differences across clinical laboratory measures, cytokines, metabolome, and gut microbiome, stratified by comparison group (IS vs. IR; DP0 vs. DP1; and subgroup comparisons such as IS-DP0 vs. IS-DP1 and IR-DP0 vs. IR-DP1). Bars indicate the number of biomarkers that show significant group differences based on Kruskal-Wallis tests, with multiple testing correction using BH (*Padj* < 0.2). This highlights domain-specific patterns in the extent to which insulin resistance and dietary pattern (independently and jointly) are associated with molecular and clinical biomarkers. **d,** Distribution of 10-year ASCVD risk scores in the study cohort. A horizontal violin plot shows the overall distribution of 10-year ASCVD risk scores, calculated using the American Heart Association Pooled Cohort Equations. **e,** Top predictors of 10-year ASCVD risk from Random Forest modeling. Bar plot shows the leading predictors identified by the Random Forest model, which incorporated demographics, dietary factors, multi-omics data, clinical lab measures, and cytokines. Feature importance values represent the relative contribution of each predictor to model performance. Bars are color-coded based on partial dependence-derived directionality: red indicates higher predicted ASCVD risk (adverse), green indicates lower predicted risk (protective), and blue indicates non-monotonic or mixed responses. PDGFBB, Platelet-derived growth factor-BB; RDW, red blood cell distribution width.

Associations between clinical lab markers, insulin resistance status, and diet patterns are shown in **Figure 5B**. In our mixed-effect models adjusting for age, sex, race/ethnicity, and baseline BMI, IR individuals showed consistently more adverse lipid profiles than IS individuals, such as higher LDL, LDL/HDL ratio, and cholesterol/HDL ratio, and lower HDL. Also, IR individuals showed elevated blood immune cell profiles, including significantly higher neutrophil counts than IS individuals. In addition, dietary pattern and insulin resistance interacted significantly on lipid markers. Specifically for LDL/HDL ratio, Tukey post-hoc comparisons revealed that IR-DP0 and IR-DP1 both had higher LDL/HDL ratios than IS-DP1 (p=0.0028 and p=0.0011, respectively), and that IS-DP1 had a lower ratio than IS-DP0 (p=0.0276). These findings suggest that diet pattern modifies lipid profiles differently in IR and IS individuals, although cautious interpretation is needed given small subgroup sizes.

### Integrative prediction models for ASCVD risk

Atherosclerotic cardiovascular disease (ASCVD) is the major cause of death and has been associated with diabetes and metabolic dysfunction. We previously reported the American Heart Association’s ASCVD risk scores of this cohort that estimate a 10-year risk of heart disease or stroke^25^. In this analysis, we characterized the distribution of ASCVD risk among study participants (**Figure 5D**). Then, we developed integrated prediction models for the 10-year ASCVD risk utilizing a comprehensive multi-domain dataset, including demographics, diet, microbiome, metabolomics, and clinical lab data **(Methods)**. We employed both Elastic Net and Random Forest prediction models within a nested cross-validation framework, calculating root mean square errors (RMSEs) to compare their accuracy and performance. This robust technique minimizes evaluation bias by using independent outer loops for performance assessments and inner loops for hyperparameter optimization, thereby preventing overfitting and enhancing model generalizability (i.e., broadly applicable results). The Random Forest models, with a 5-fold cross-validation and 500 trees, outperformed the Elastic Net models in predicting the ASCVD risk score. The final Random Forest model, presented in **Figure 5E**, highlights the top 15 most influential features (calculated RF importance). Additionally, partial dependence plots (PDPs) were used to identify the direction of these features’ impacts (positive or negative or mixed) on the ASCVD risk score.

Age emerged as the strongest predictor of ASCVD risk, followed by total fiber intake, aminoadipic acid, seafood intake, platelet-derived growth factor subunit B (PDGFBB), adipylcarnitine, and specific microbial taxa (e.g., class *Bacilli*, order *Selenomonadales*, and class *Negativicutes*). ASCVD risk increased with age, particularly after 50 years old. The risk also increased with higher platelet-derived growth factor-BB (PDGFBB), red blood cell distribution width (RDW), and specific metabolites and microbial taxa. In contrast, higher total fiber intake was associated with a lower risk, showing a steep drop and then reaching a plateau at ∼ 10 grams. Seafood consumption was generally protective, although very high seafood intake (> 300 kcal per 1,000 kcal) may increase the risk. Copper intake showed a pronounced protective effect against CVD risk.

## Discussion

Despite extensive research on the link between diet and metabolic health, fundamental questions remain underexplored regarding how longitudinal changes in habitual dietary patterns dynamically influence the gut microbiome and host metabolome over time, and whether an individual’s metabolic phenotype, specifically insulin resistance status, modifies these relationships. Through year-long, multi-omics profiling of 71 deeply phenotyped adults, we addressed these gaps using gold-standard insulin suppression tests to determine insulin resistance status, machine learning-derived dietary patterns, and a novel multidimensional feature-based analytical approach that captures both magnitude and temporal complexity of diet-omics interactions.

Our findings reveal two main advances with direct implications for precision nutrition. First, we demonstrate that longitudinal diet-omics interactions are modified by insulin resistance status. Using a temporal shape feature approach that characterizes both the magnitude and patterns of change in nutrient intake over time, we found that insulin-sensitive (IS) individuals exhibited significantly more numerous and stronger correlations between dietary changes and both metabolomic and microbial shifts compared to insulin-resistant (IR) individuals. IS individuals showed pathway enrichment in steroid hormone, sphingolipid, pyrimidine, and phenylalanine metabolisms, as well as higher density of nutrient-metabolite correlations within gamma-glutamyl amino acid, branched-chain amino acid, butanoate, vitamin D, purine, and acyl-glycine-related fatty acid metabolisms.

This pattern of differential responsiveness, largely absent in IR individuals, positions insulin resistance as a state of reduced molecular adaptability to dietary changes. The attenuated diet-omics coupling observed in IR individuals is consistent with impaired metabolic flexibility and altered substrate utilization characteristic of the insulin-resistant phenotype. Although these mechanisms were not directly measured in our study, the observed reduction in diet-responsive molecular networks suggests that baseline metabolic state shapes the extent to which dietary changes translate into coordinated metabolomic and microbial remodeling. Notably, recent evidence shows that diet and exercise interventions differentially modulated microbiome-associated metabolites, including those related to impaired glucose regulation. This supports the concept that baseline metabolic health influences molecular responsiveness to lifestyle modification^16^. Together, these findings identify insulin resistance status as a critical modifier of diet-induced molecular responses, highlighting metabolic health as an important determinant in precision nutrition. Whether restoration of metabolic flexibility enhances responsiveness to dietary intervention remains an important question for future interventional studies.

Second, we integrated diet, multi-omics profiles, clinical laboratories, inflammatory biomarkers, and demographic characteristics into a machine learning-based model to predict 10-year atherosclerotic cardiovascular disease (ASCVD) risk. Beyond traditional risk factors, specific metabolites and microbial features emerged as prominent predictors. Several top ranked metabolites were consistent with pathways related to cellular energy handling and substrate oxidation^42,43^. Alpha aminoadipic acid (2-AAA), one of the strongest predictors, has previously been associated with increased heart disease risk, visceral adiposity and hepatic steatosis^43^, and has been proposed as a marker of disrupted lysine catabolism and metabolic stress. Elevated adipylcarnitine levels similarly aligned with a recent meta-analysis of 416 studies^44^ that identified many middle and long-chain acylcarnitines among the 23 prognostic metabolic markers for cardiovascular events. Accumulation of these intermediates is consistent with incomplete fatty acid oxidation and perturbed metabolic flux, processes increasingly implicated in cardiometabolic dysfunction. While our analysis does not establish causality, it highlights coordinated alterations in energy metabolism as components of cardiovascular risk.

The gut microbiome features also contributed to model performance, suggesting that host metabolic state and gut microbial ecology jointly inform cardiovascular risk. Class Bacilli, such as Streptococcus, has been linked to atheromatous plaques^45^, supporting a potential interface between microbial composition and vascular pathology. Other taxa, Class Negativicutes and Order Selenomonadales, may influence CVD risk through less direct pathways, potentially via sleep deprivation as suggested by recent Mendelian randomization studies^46^. Future research integrating wearable sleep monitoring with microbiome profiling could further elucidate these connections.

Fiber intake exhibited a non-linear association with predicted cardiovascular risk, with risk reduction appearing to plateau at approximately 10 grams per 1,000 kcals in this cohort. Although dietary guidelines recommend 14 grams of fiber per 1,000 kcals (IOM), our findings suggest minimal additional benefit beyond this level. This pattern is consistent with established cardioprotective effects of fiber, including modulation of lipid metabolism, improved postprandial glycemic control, bile acid regulation, and microbial production of short chain fatty acids. Higher intakes may confer additional benefits across other metabolic and gastrointestinal outcomes. More broadly, several predictors, including seafood consumption, demonstrated non-linear relationships with cardiovascular risk, highlighting the complex, multifactorial nature of diet-related cardiovascular biology.

To evaluate whether gut microbial features bridge dietary components (i.e., nutrients, food groups, and deep-learning identified dietary pattern) and host metabolic profiles, we conducted mediation analyses integrating dietary exposures, genus-level microbiota, and plasma metabolites. Although mediation effects did not survive correction for multiple testing in this modestly sized cohort (n = 71), *Parabacteroides* emerged across models of a refined carbohydrate-rich dietary pattern with both N-acetyl-1-methylhistidine (indirect effect = -0.12), an amino acid-derived metabolite associated with kidney function^47,48^, and PE(P-36:4) (indirect effect = -0.09), a plasmalogen phosphatidylethanolamine involved in redox-sensitive lipid remodeling^49–51^. Protein intake did not differ across dietary pattern groups, indicating that these associations are unlikely to reflect variation in precursor availability alone. This finding aligns with recent evidence that diet-induced changes in phosphatidylethanolamine species are among the most informative lipidome features between dietary fat quality and cardiovascular and type 2 diabetes risk reduction^52^. Instead, the convergence of dietary pattern, microbial abundance, and metabolite signatures suggests coordinated restructuring of host-microbial metabolic networks. Although the direct mechanism connecting *Parabacteroides* to these metabolites is not established, prior multi-omics studies indicate that gut microbes can influence host phospholipid and amino acid metabolism through bile acid signaling, inflammatory modulation, and oxidative stress pathways^52,53^. The recurrence of *Parabacteroides* across distinct metabolic domains identifies the microbiome as a candidate intermediary in diet-associated metabolic remodeling. Clarifying whether specific taxa causally mediate dietary effects on host metabolism will require larger longitudinal and interventional studies.

This study has several limitations. First, while the longitudinal design with dense sampling enhances our capacity to detect dynamic associations, our findings primarily establish associations and statistical mediation effects rather than direct causal relationships. Due to the observational nature of the study, causal inference cannot be drawn with certainty. Additionally, some effects with small samples may not have been detected. Future intervention trials with larger sample sizes could confirm causal relationships. Second, because our cohort was primarily drawn from the San Francisco Bay Area, the generalizability of our findings is limited to populations with similar demographics and geographical characteristics. However, recent studies have shown that diet-metabolite associations are reproducible across ethnically diverse populations^54^ and that microbiome-metabolite signatures are consistent across multiple independent cohorts^16^. Third, at the time of study initiation, 16S rRNA sequencing was an efficient method for analyzing microbiomes rich in human DNA, but it has limitations in accurately detecting specific bacterial genera^27^. Metagenomic sequencing could provide a more comprehensive view in future studies.

In conclusion, this study provides a comprehensive, longitudinal multi-omics perspective on how habitual diet shapes host metabolism and the gut microbial ecology across different metabolic phenotypes (IS vs IR). We show that insulin resistance dampens diet-associated molecular remodeling across both metabolomic and microbial domains, identifying metabolic flexibility as a central feature influencing dietary responsiveness. *Parabacteroides* emerged as a potential microbial mediator linking dietary patterns to host metabolic profiles. Importantly, our integrative cardiovascular risk model showed that diet, plasma metabolites, gut microbial taxa, and immune markers each contributed to ASCVD risk. These findings support a systems perspective in which cardiometabolic disease reflects interactions between host metabolism and the gut microbiome. By combining longitudinal dietary assessment with multi-domain molecular profiling, this work moves beyond single nutrient approaches and advances a biologically informed framework for precision nutrition. Together, our results suggest that differences in metabolic responsiveness to diet may contribute to variation in cardiometabolic risk, with implications for targeted prevention strategies.

## METHODS

### Cohort Description and Study Design

Our research group previously established a cohort comprising more than 100 adults at risk of type 2 diabetes (T2D) from the San Francisco Bay Area in California through advertisements in local newspapers and radio broadcasts. The participants were then followed longitudinally from 2013 to 2017. Comprehensive details of this study design are previously described elsewhere^25–28^. Briefly, exclusion criteria included anemia (hematocrit levels <30), renal disease (creatinine levels >1.5), a history of cardiovascular diseases, malignancy, chronic inflammatory or psychiatric conditions, or prior bariatric surgery or liposuction. Participants provided blood and stool samples following a 12-hour fasting every three months and completed food diaries at Stanford’s Clinical and Translational Research Unit (CTRU). All participants provided written informed consent under research protocol 23602, approved by Stanford University’s Institutional Review Board. Deep phenotyping of participants was performed through longitudinal multi-omics profiling of biospecimens collected at each study visit across the cohort. Plasma samples were analyzed to quantify metabolites, cytokines, and clinical laboratory markers and stool samples were used to assess the gut microbiota. This approach enabled a comprehensive, longitudinal analysis of both systemic and gut-specific biological features in the study population.

### Food Diary and Diet Data Analysis

Participants were instructed to choose a typical weekday from the past week, that was most representative of their eating habits over the past three months. They were asked to record all of their food consumption, including condiments, and the methods of food preparation such as baking, frying, roasting, breading, etc. Additionally, they were encouraged to provide homemade recipes that were used to prepare any food that was consumed. Under the dietitian’s guidance, participants were urged to be as precise as possible in detailing the type, portion size, brand name, eating time, time of food consumption, and even the name of the restaurant, if applicable. They were given examples to guide their records. For example, “8 AM: two slices of Alpine Valley whole wheat bread, 1 slice of Finlandia Swiss cheese, half of one medium sized avocado, 1 large hardboiled egg, and 1 cup of Kirkland Signature non-fat milk.” They were instructed to document all their meals of the day: breakfast, lunch, dinner, snacks, and any other information relevant to their diet.

Food logs collected from study participants were analyzed for dietary composition (nutrients and food groups) by research dietitians using the ESHA Food Processor Nutrition Analysis software (ESHA Research, Salem, OR). The prime data sources were the USDA-SR-28 and the USDA’s Food and Nutrient Database for Dietary Studies (FNDDS) on the Food Processor. If participants did not specify quantities, we referred to the corresponding gender and age data from the Diet History Questionnaire III (DHQ III) and selected a medium portion size, as the DHQ III is based on a compilation of national 24-hour dietary recall data from the National Health and Nutrition Examination Surveys (NHANES). Additionally, if participants reported that they consumed different foods regularly, such as having either eggs or cereal for breakfast, then both items were recorded but each was entered as half the amount to reflect typical intake and ensure the most accurate nutrient profile. The study evaluated 62 nutrients, which included energy, carbohydrate, protein, fat, fiber, etc. Additionally, 20 food groups were assessed, such as Bread Products, Cereals & Grains, Coffee, Dairy Products, Eggs, Fats & Oils, Fruits, Legume Products, Meat & Poultry, Pizza & Sandwich & Wrap, Non-Starchy Vegetables, Nuts & Seeds, Rice & Pasta & Noodles, Salad, Starchy Vegetables, Sweets & Sweeteners, Sauces & Spices & Herbs, Snacks, Seafoods, and Tea. Once dietary data were collected, they were matched with corresponding omics data collection dates to investigate potential relationships between changes in diet and omics profiles. 87% of the dietary records were matched with omics data within a 2-week window, while the remaining records were matched within a window of up to three months.

### Insulin Suppression Test (Determination of Insulin Sensitivity)

A subset of eligible consenting participants (n=45) underwent a one time insulin suppression test (IST) to determine their insulin sensitivity by assessing insulin-mediated glucose uptake^25,26^. Following a 12-hour overnight fast, participants received an intravenous infusion consisting of 0.27 ug/m2 min of octreotide, 25 mU/m2 min of insulin, and 240 mg/m2 min of glucose over a period of 180 minutes at CTRU. Blood samples were collected every 10 minutes during the last 30 minutes (150-180 min) of the infusion, totaling four blood draws. Plasma glucose concentrations were measured using an oximetric method, while insulin levels were quantified by radioimmunoassay. The mean value of the four steady-state plasma glucose (SSPG) and insulin concentrations were calculated. Participants were subsequently categorized based on their SSPG results: insulin sensitive (IS) (n=18) if SSPG < 150 mg/dl, or insulin resistant (IR) (n=27) if SSPG ≥ 150 mg/dl. Data were available only for a subset of participants due to several reasons, including withdrawal from the study before their scheduled tests, opting out due to scheduling conflicts or personal preferences; or lacking dietary data (n= 26).

### Untargeted Metabolomics Analyses

Participants’ plasma samples were collected in EDTA tubes and processed within 24 hours. Metabolites were extracted from plasma aliquots using a 1:1:1 solvent mixture of acetone, acetonitrile, and methanol, followed by evaporation under nitrogen and reconstitution in a 1:1 methanol:water solution. Untargeted metabolomics profiling was conducted using liquid chromatography-mass spectrometry (LC-MS) with both hydrophilic interaction liquid chromatography (HILIC) and reverse-phase liquid chromatography (RPLC) separations, under positive and negative electrospray ionization modes. Data were acquired on a Thermo Q Exactive Plus mass spectrometer equipped with a HESI-II probe, and MS/MS spectra were collected from pooled study samples consisting of an equimolar mixture of 100 randomized samples.

Quality control procedures were implemented, including sample randomization for both metabolite extraction and data acquisition, use of 9 isotopically labeled internal standards during sample preparation to monitor extraction efficiency and LC-MS performance, and regular injections of pooled samples throughout the run to assess signal stability.

Data processing was performed using Progenesis QI software (v2.3, Nonlinear Dynamics, Waters), with feature filtering based on dilution linearity and presence in at least 33% of samples. Signal drift was corrected using LOESS normalization.

Metabolites were annotated using MS/MS fragmentation spectra and categorized based on the Metabolomics Standards Initiative (MSI) confidence levels. The final metabolite results are classified into pathways using KEGG and HMDB annotations and retained for downstream analysis. Further methodological details are found elsewhere^25–28^

### Quantification of Clinical and Inflammatory Markers

Clinical lab tests were conducted at the Stanford Clinical Laboratories. The standardized procedures for collecting and submitting blood and urine biospecimen can be found elsewhere (https://stanfordlab.com/test-directory.html). These tests include a metabolic panel, complete blood count panel, glucose, hemoglobin A1c (A1C), insulin, high-sensitivity C-reactive protein, immunoglobulin M, and lipid, kidney, and liver panels.

Plasma cytokines, chemokines, and growth factors were measured, following established procedures from the Stanford Human Immune Monitoring Center (HIMC). The Human 62-plex Luminex multiplex assay using conjugated antibodies (Affymetrix, Santa Clara, California) was performed. Specifically, the raw data from the assay were normalized against the median fluorescence intensity value, and then, a variance stabilizing transformation was applied to eliminate the batch effects. Outliers that had background noise (CHEX) > 5 standard deviations from the mean were excluded from the dataset.

### Microbiome Analysis (16S Microbiome Sequencing)

Stool samples were stored at -80 °C immediately after arrival. Stool DNA was extracted following the Human Microbiome Project’s Core Sampling Protocol A (https://www.hmpdacc.org). The V1-V3 hypervariable regions of the 16S rRNA gene were amplified using the primer pair 27F (5′-AGAGTTTGATCCTGGCTCAG-3′) and 534R (5′-ATTACCGCGGCTGCTGG-3′). Amplicons were then sequenced using paired-end 2×300 base-pair reads on the Illumina MiSeq platform. Initial base calling and demultiplexing were performed using Illumina’s software, allowing one mismatch in the primer sequence and no mismatches in sample-specific barcodes. Barcode and primer sequences were removed before downstream analysis. Reads were filtered to exclude sequences with ambiguous bases or low quality (Q-score <35). High-quality reads were clustered into operational taxonomic units (OTUs) using the Usearch algorithm, and taxonomic assignments were made using the RDP Classifier against the Greengenes reference database (May 2013 version). Detailed methods are described previously^25,26^.

### ASCVD Risk Score Calculation

The 10-year ASCVD risk was calculated using the 2013 ACC/AHA Pooled Cohort Equations^55^, following guideline instructions, with analyses performed in SAS version 9.4. Although newer tools such as the 2023 AHA PREVENT equations have been released, the original PCE equations continue to be widely used in analyses up to and including those referenced in the 2023 guidelines. For all participants, baseline time points were used and details are provided elsewhere^25^.

## Data Analyses

### Baseline characteristics and longitudinal dietary analyses in the overall cohort and by insulin resistance status

To compare baseline demographics, laboratory measures, and metabolic test results between insulin-sensitive (IS) and insulin-resistant (IR) participants, we used the Wilcoxon rank-sum test for continuous variables not normally distributed, and the χ² test or Fisher’s exact test for categorical variables, as appropriate.

Longitudinal changes in nutrient and food group intake in the overall cohort and within IS and IR groups were evaluated using linear mixed-effects models. Nutrient and food group intakes were log-transformed before modeling. Estimated marginal means were used to quantify changes from baseline (T1) and to generate predicted means on the original scale. P values for within-group time contrasts were adjusted using the Benjamini-Hochberg (BH) procedure. To compare nutrient/food group intakes between IS and IR participants at each time point and to compare intakes between DP0 and DP1, we used Mann-Whitney rank-sum tests with BH-adjusted P values.

### Unsupervised machine learning of dietary patterns

A deep autoencoder framework was used to extract latent features from the food log data using keras with a TensorFlow backend. Six input matrices were analyzed: two datasets on energy intake proportion from each food group, two on food weight from each food group, and two on carbohydrate intake proportion from each food group. To prevent model overfitting and retain only reproducible feature representations, we performed an extensive model search. Specifically, 19 different autoencoder architectures, varying in depth, width, and regularization settings, were each trained with 10 independent random seeds for weight initializations. As a result, we obtained 1,140 initial autoencoder trained models. Next, the models were retained only if they demonstrate both accurate and stable performance. Retention criteria included a mean validation reconstruction loss < 0.005 across 5-fold cross-validation, and a coefficient of variation for validation loss < 20%. This ensured both minimal reconstruction error and high consistency across folds. Finally, k-means clustering was performed on the extracted features from the remaining models (n=650) with the “scikit-learn” package. We selected two clusters (k = 2), which yielded high silhouette scores (> 0.35) and showed a standard deviation of cluster size proportion (< 0.15). These criteria ensured both compactness and stability of clusters. We also visually confirmed the separation of clusters using both principal component analysis (PCA) and t-distributed stochastic neighbor embedding (t-SNE) plots (**Figure 2C**).

### Cross-sectional analysis of diet-omics associations in the study cohort

We used partial Spearman correlation analyses, adjusting for age and sex, to examine associations between each dietary variable (62 nutrients and 20 food groups and a binary ML-dietary pattern) and each omics feature (724 plasma metabolites and 109 microbiota taxa across multiple taxonomic levels). Prior to analysis, multiple metabolome measurements of each participant were aggregated to the median value. The dietary pattern variable contrasted individuals classified as DP1 (refined carbohydrate richer pattern) versus DP0, with DP0 serving as the reference group. Multiple testing was addressed using the Benjamini-Hochberg procedure, with significance defined as a BH-adjusted p value < 0.20. Results of both metabolome and microbiome were visualized using a clustered heatmap with dietary components. To summarize the overall pattern of diet-metabolome associations, we created a Sankey diagram which connects nutrient categories (carbohydrates, protein, fat, minerals, fat-soluble and water-soluble vitamins, fiber, and caffeine) to the top 10 metabolic pathways based on the number of significant correlations (*P* < 0.01).

### Mediation analysis (Diet to Omics)

We conducted mediation analyses to investigate whether gut microbiome composition mediates the association between dietary components (nutrients, food groups, and ML-derived binary dietary pattern) and plasma metabolites. Given the large number of dietary exposures, including 62 nutrients, 20 food groups, and binary dietary pattern, mediation analyses were limited to diet-metabolite associations that showed statistical significance in prior cross-sectional models in order to reduce the multiple testing burden and improve computational feasibility and model stability. For each association, mediation models were specified to test the indirect effect of diet on metabolites through genus level microbiome taxa, while accounting for the direct effect of diet. Linear regression-based mediation models were applied using log-transformed microbiome and metabolite data, with age and sex adjustment. Indirect effects were estimated as the product of the diet to microbiome and microbiome to metabolite pathways, and statistical significance was evaluated. Multiple testing across mediation models was explored using the Benjamini-Hochberg procedure. Diet-microbiome-metabolite relationships with significant indirect effects (unadjusted *P* < 0.05) were interpreted as microbiome-mediated associations between dietary components and circulating metabolites.

### Correlation network analysis

To explore longitudinal relationships among nutrients, gut microbiota, and plasma metabolites, we aligned nutrient intake, gut microbiome, and metabolomics data within individuals, ensuring that sample collection dates were within one week of each other. Then, we derived 11 temporal change features for each analyte, summarizing longitudinal dynamics through the study (e.g., early and late slopes, cumulative exposure, and peak change). Correlation networks were then constructed separately for insulin-sensitive (IS) and insulin-resistant (IR) groups.

### Nutrient-microbiome correlation network

For each group (IR and IS), Spearman correlation coefficients were calculated for all nutrient-microbe feature pairs. Only associations with FDR q value < 0.05 were retained for network construction.

### Nutrient-metabolite correlation network

Similarly, Spearman correlations were computed between nutrient-metabolite feature pairs within each group, and only statistically significant correlations (FDR q < 0.05) were included in the final network and used for pathway-level density comparisons.

### Metabolic pathway comparison between IR and IS

To compare pathway-level metabolic responses between IR and IS individuals, we conducted a bootstrapped pathway-level analysis based on the significant nutrient-metabolite correlation results (FDR q<0.05) using pre-computed correlation datasets. For each group, we quantified the sub-pathway density by calculating two metrics: (1) the total number of significant nutrient-metabolite associations mapped to a given sub-pathway, and (2) the number of unique metabolite-nutrient pairs contributing to that sub-pathway. The ratio of these two values was used as a normalized density score for each sub-pathway within each group.

To assess statistical differences between groups, we performed 1,000 bootstrap iterations. In each iteration, we resampled the correlation dataset with replacement, recalculated the enrichment density scores for all sub-pathways in both groups, and aggregated the results. We then used the Wilcoxon rank-sum test to evaluate the significance of density differences between IR and IS groups for each sub-pathway. Sub-pathways with significant differences (P < 0.05) were visualized using bar plots displaying group means and standard deviations of bootstrapped density scores. The top 10 sub-pathways with the largest absolute differences between groups were highlighted to showcase the most divergent metabolic pathways.

### Simultaneous and lagged correlation analysis between nutrient intake and gut microbiota

To investigate the dynamic relationship between nutrient intake and gut microbial composition, we performed both simultaneous (synchronous) and lagged correlation analyses using longitudinal data from participants with complete data across all four time points. We aligned nutrient intake, gut microbiome, and metabolomics data within individuals, ensuring that sample collection dates were within one week of each other.

### Simultaneous correlation analysis

For each individual, we computed the difference in nutrient intake and gut microbiota abundance between paired time points to capture temporal changes. Specifically, we derived six nutrient-microbiome “features” representing differential pairs (denoted as Features 5-10; **Figure 4A**). For each feature, we then calculated the Pearson correlation coefficient between nutrient intake and microbial abundance across individuals. Based on the time interval between paired time points, we classified these correlations into three categories: short-term correlations (3-month intervals; features 5, 9, and 10), medium-term correlations (6-month intervals; features 6 and 8), and long-term correlations (9-month interval; feature 7).

### Lagged correlation analysis

In parallel with the simultaneous correlation analysis, we also examined lagged correlations between nutrient intake and gut microbiota. We first calculated changes in nutrient intake and microbial abundance across different time points, as described above. We then assessed the temporal lag between nutrient exposure and subsequent microbiome response by computing Pearson correlations between non-concurrent features: (1) Nutrient data from feature 5 and microbiome data from feature 6. (2) Nutrient data from feature 5 and microbiome data from feature 7. (3) Nutrient data from feature 6 and microbiome data from feature 7. (4) Nutrient data from feature 9 and microbiome data from feature 8. Based on the time intervals between these pairs, we categorized the resulting correlations as follows: (a) Short-term lagged correlations: pairs (1) and (4), corresponding to 3-month lags; (b) Medium-term lagged correlations: pair (3), corresponding to a 6-month lag; and (c) Long-term lagged correlations: pair (2), corresponding to a 9-month lag. This approach allowed us to capture delayed associations between nutrient intake and gut microbial dynamics over different temporal scales.

### Cytokine and clinical laboratory associations with insulin resistance and dietary pattern

To evaluate the independent and interactive associations of insulin resistance (IR) and dietary pattern (DP) with systemic inflammatory and metabolic markers, we analyzed 62 circulating cytokines and 43 clinical lab measures using linear mixed-effects (LME) models. Biomarker concentrations were log-transformed and modeled as the dependent variable, with fixed effects for insulin resistance (SSPG as a continuous variable for cytokine analyses; IS/IR as a categorical variable for clinical lab analyses), diet pattern (DP0, DP1), and their interaction, and with sex, age, race/ethnicity, and baseline BMI included as covariates. Participant ID was included as a random intercept. This approach allowed us to assess both main effects and interaction effects between insulin resistance and dietary pattern. Multiple testing adjustment was applied to interaction *P* using BH method (*Padj* < 0.2). For clinical laboratory markers with significant interaction terms, we conducted Tukey post-hoc pairwise comparisons among the four subgroups (IR-DP0, IR-DP1, IS-DP0, IS-DP1) using estimated marginal means from the same LME models. Cytokine interaction was visualized using regression plots stratified by dietary pattern, whereas clinical laboratory markers were visualized as a heatmap of estimated marginal means (standardized as z-score). Clinical lab markers were further grouped into physiological domains such as electrolytes, glucose regulation, hepatic function, immune, lipid, renal, and hematologic systems.

### ASCVD Risk Score Prediction Models (Elastic Net and Random Forest Models)

After predictor variables were imputed using missForest, nested cross-validation was employed, with each iteration splitting the data into training and test sets. Feature selection was performed on the training data using LASSO to identify the features to be retained. For model training, an inner cross-validation control with 5 folds was established. Both Elastic Net and Random Forest prediction models were conducted to assess and compare model accuracy and performance. Model predictions were generated for the respective outer test sets, and the root mean square error (RMSE) was calculated for each model. After completing all iterations of all outer folds, the mean RMSEs for each model were compared using a t-test.

Final models were trained using the tuning parameters derived from nested cross-validation on the full dataset. The LASSO model was fitted using the optimal lambda value, and features were selected accordingly. Only these selected features were used to train the final Elastic Net model, which was optimized through a grid search of alpha and lambda values for hyperparameter tuning. Coefficients from the final Elastic Net model were extracted, and feature importance was assessed based on the absolute values of these coefficients. The top 20 most significant features were then chosen for visualization.

Similarly, the final Random Forest model was trained using 5-fold cross-validation with 500 trees. Feature importance scores were extracted from this model, and the top 20 features were also selected for visualization. To determine the direction and magnitude of each feature’s influence on ASCVD risk, Partial Dependence Plots (PDPs) were generated. Based on the RMSE comparisons, the Random Forest model and Elastic Net models showed similar performance, but RF had a slightly lower RMSE. Therefore, we present the results from the Random Forest model as the main figure in our study.

## Acknowledgements

We thank all study participants and investigators for their contributions. We acknowledge Tanya Chettri, Keya Mann, Leah Jones, Syed Shams, and Agathe Grisard for assistance with dietary data analysis under the supervision of our study registered dietitians. We thank Lettie McGuire for preparation of the graphical abstract. This work was supported by the National Institutes of Health (NIH) grants U54DK102556, R01DK110186, R01HG008164, S10OD020141, UL1TR001085 and P30DK116074 (to M.P.S.), and by the Stanford Data Science Initiative (to M.P.S.). H.P. was supported by NIH institutional training grant T32HL098049 and by the Stanford Lifestyle Medicine Program. SMS-FR was supported by NIH grant K08 ES028825. The funders had no role in study design; data collection, analysis or interpretation; manuscript preparation; or the decision to submit the manuscript for publication.

## Data Availability

The datasets generated and analyzed during this study will be available from the corresponding author upon reasonable request. Deidentified, non-protected health information data will be deposited in a publicly accessible repository at the time of publication.

## Code Availability

All code used for data processing and analysis will be made publicly available upon publication at https://github.com/heyjunp/ipop_diet_omics.

## Author Contributions

H.P. and M.P.S. conceived and designed the study. H.P. and X.S. developed the analytical framework, performed the primary analyses, and generated figures. P.B. and Y.L. conducted additional analyses and contributed to figure preparation. H.P., D.P. and R.B. supervised and performed dietary data processing and analyses. A.C. and C.B. coordinated study operations. H.P. drafted and H.P. and M.P.S. revised the manuscript, with input from all authors. All authors reviewed and approved the final manuscript.

